## Supplementary figures and images for "The requirement of GW182 in miRNA-mediated gene silencing in *Drosophila* larval development"

### Figures EV1-EV-5

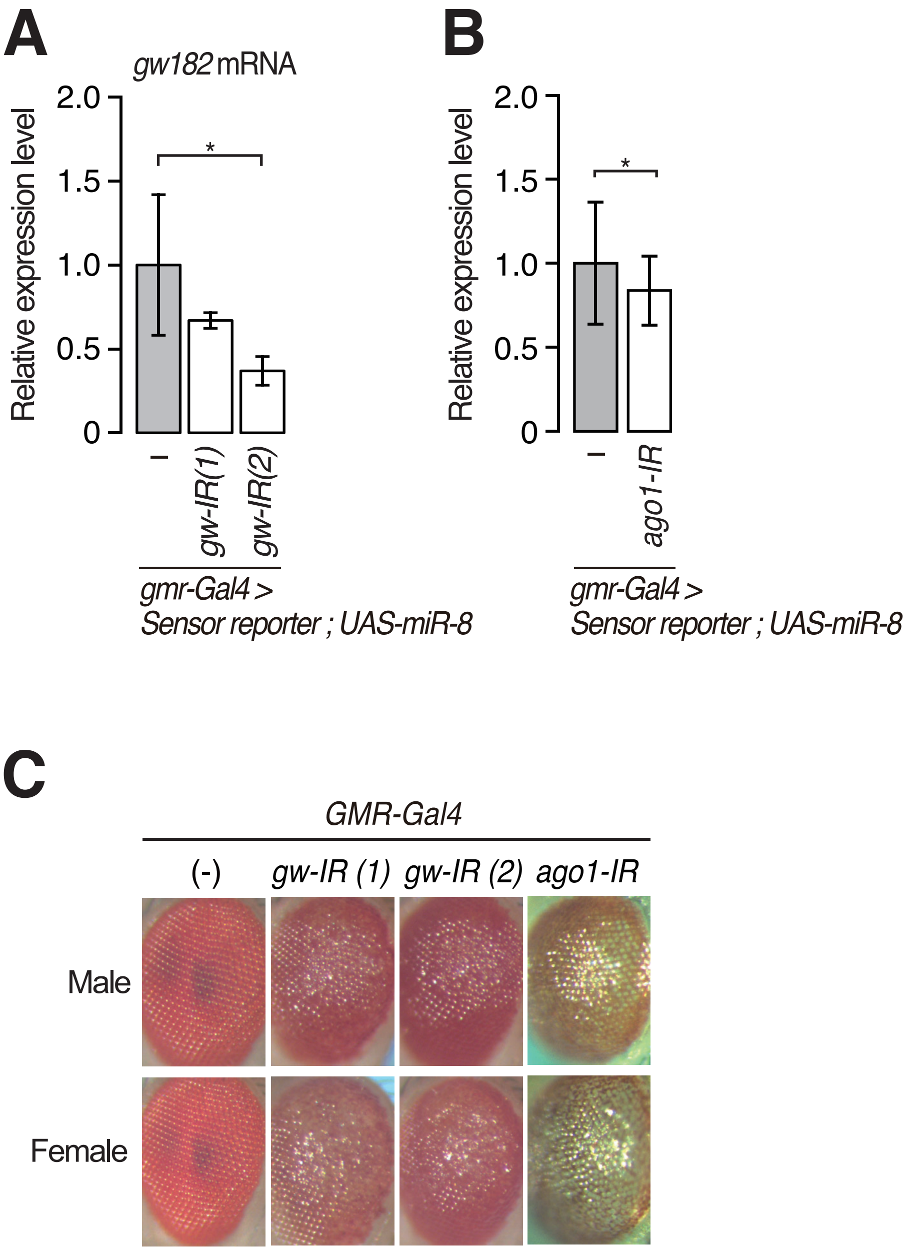

#### Figure EV2

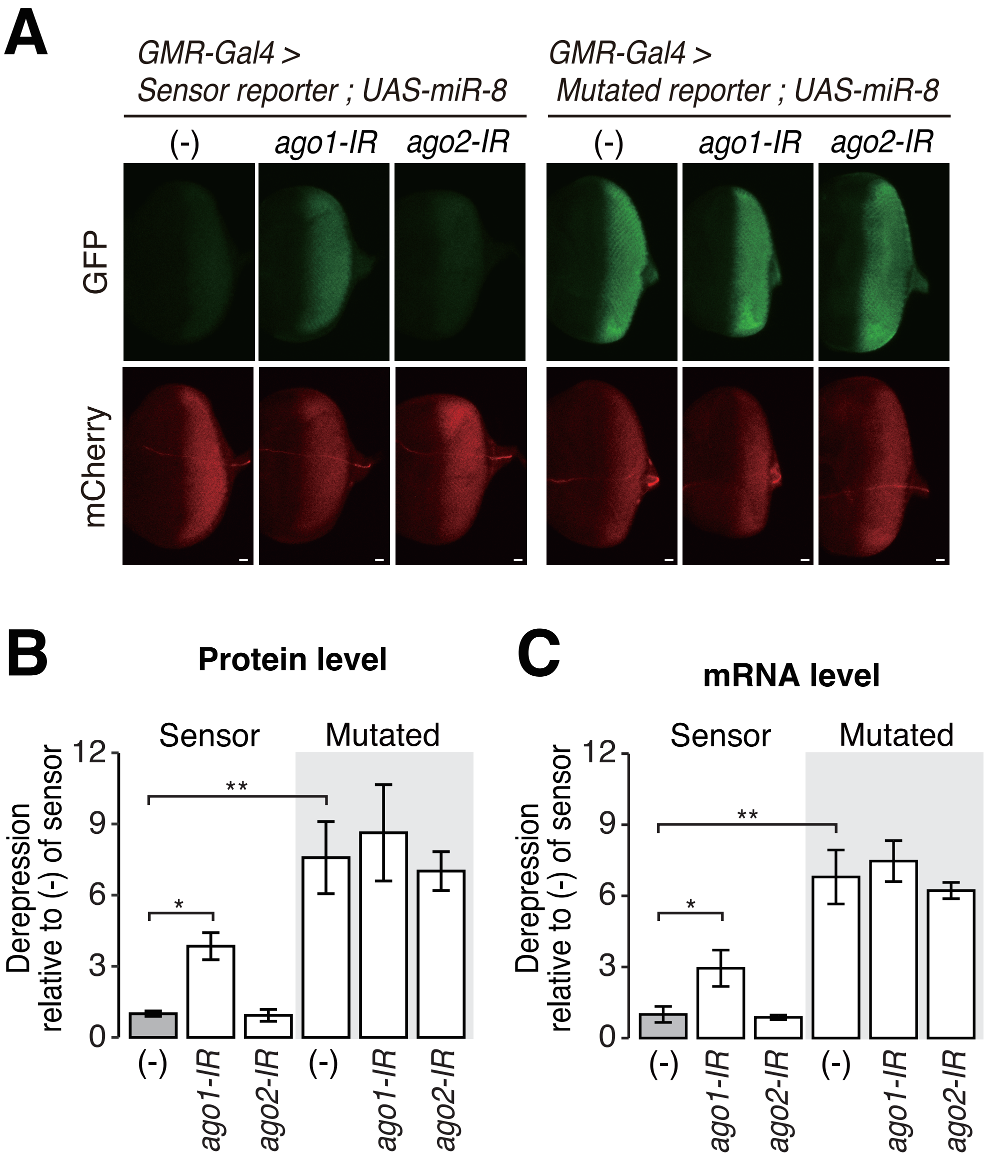


Matsuura-Suzuki et al.

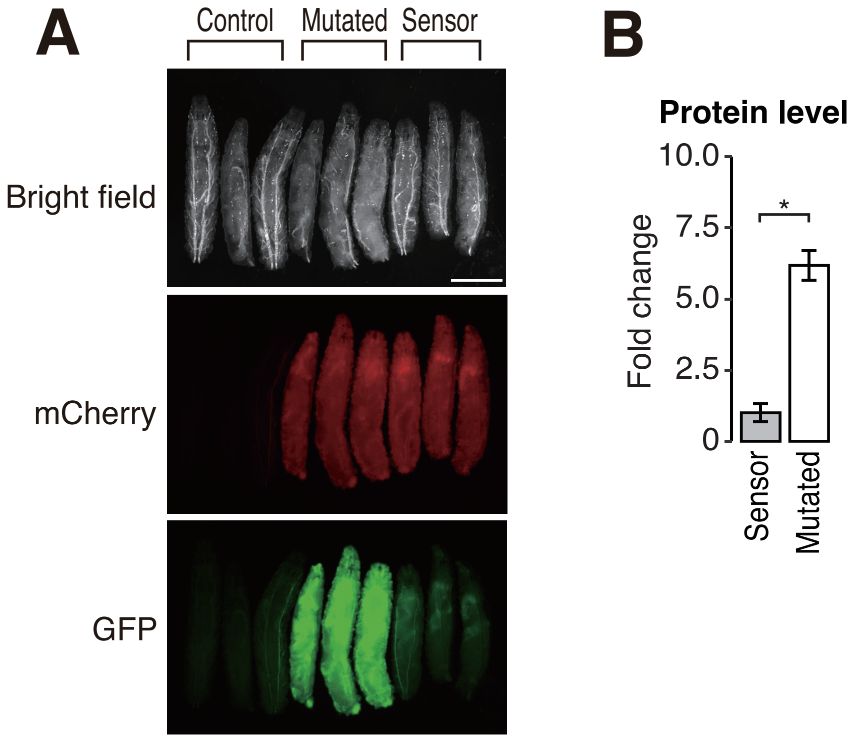

#### Figure EV4

**A**

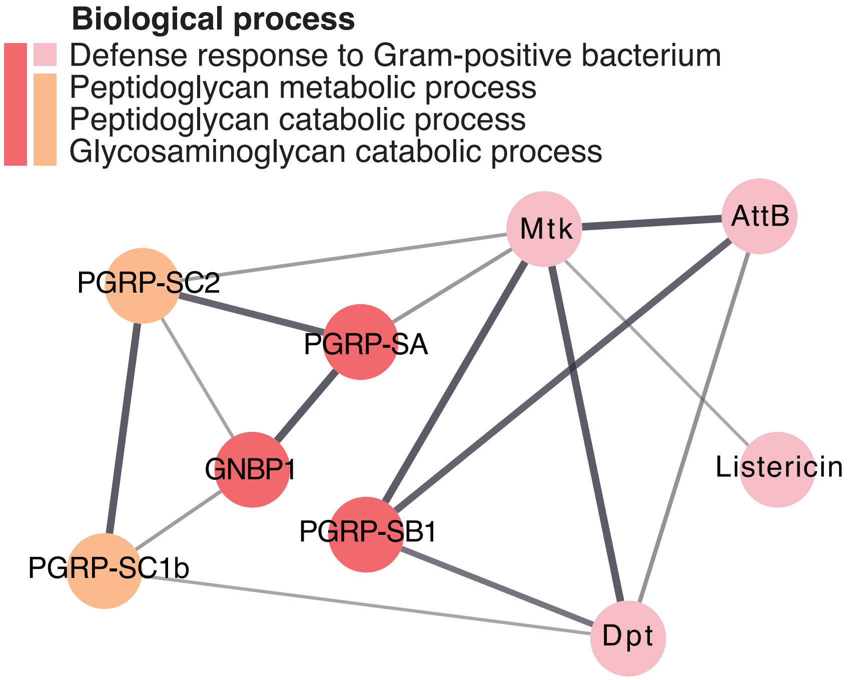


**B**

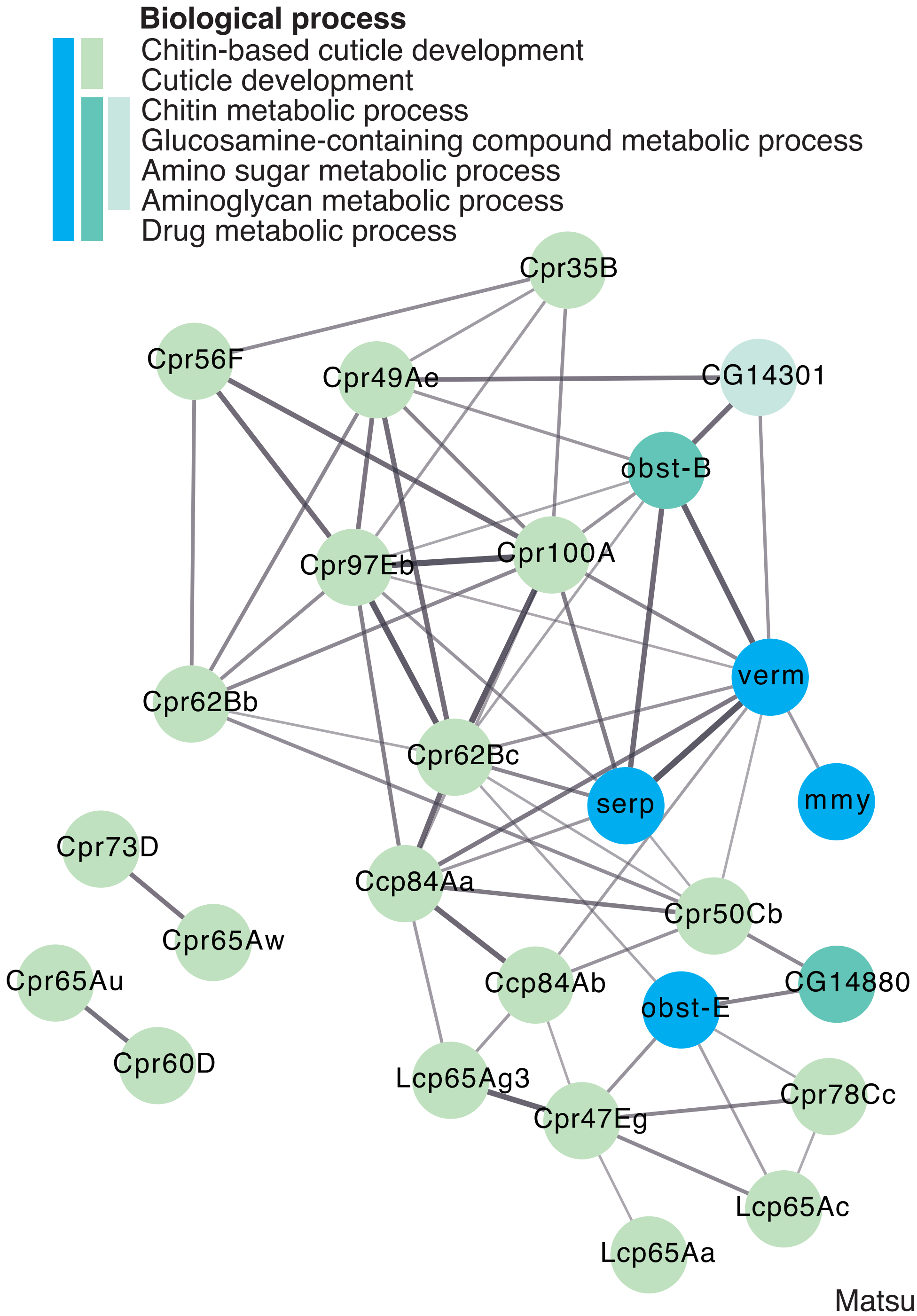


Matsuura-Suzuki et al.

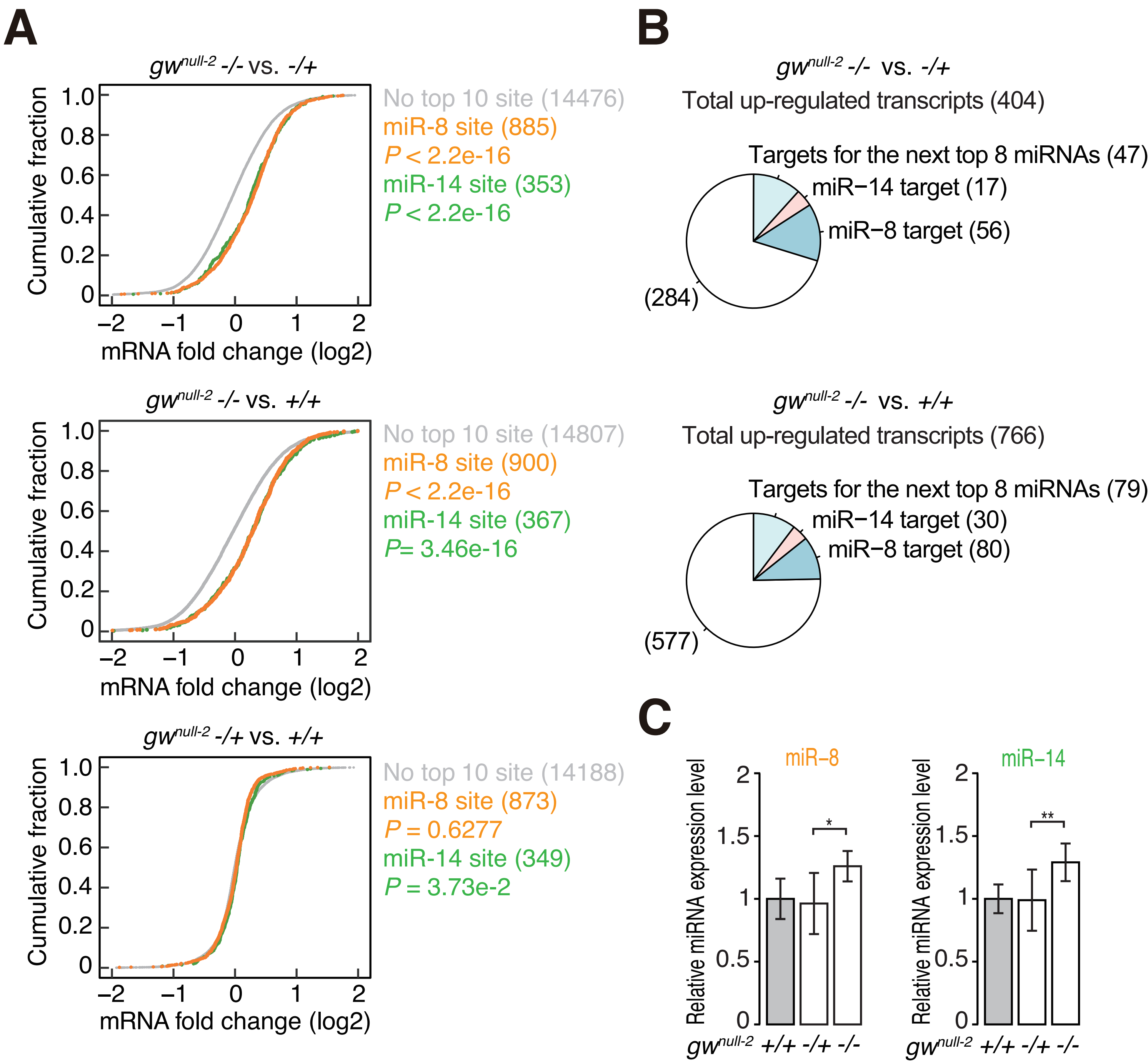
